# Niclosamide targets macrophages to rescue the disrupted peritoneal homeostasis in endometriosis

**DOI:** 10.1101/2022.04.05.487220

**Authors:** Liang Zhao, Mingxin Shi, Sarayut Winuthayanon, James A. MacLean, Kanako Hayashi

**Author notes:** Correspondence to Kanako Hayashi. Disclosures: The authors declare no competing interests exist.

## Abstract

Due to the vital roles of macrophages in the pathogenesis of endometriosis, targeting macrophages could be a new therapeutic direction. Here, we investigated the efficacy of niclosamide for the resolution of perturbed microenvironment caused by dysregulated macrophages in a mouse model of endometriosis. Single-cell transcriptomic analysis revealed the heterogeneity of macrophage subpopulations including three newly identified intermediate subtypes with sharing characteristics of traditional “small” or “large” peritoneal macrophages (SPMs and LPMs) in the peritoneal cavity. Endometriosis-like lesions (ELL) enhanced the differentiation of recruited macrophages, promoted the replenishment of resident LPMs, and increased ablation of embryo-derived LPMs, which were stepwise suppressed by niclosamide. In addition, niclosamide reversed intercellular communications between macrophages and B cells which were disrupted by ELL. Therefore, niclosamide rescued the perturbed microenvironment in endometriosis through its fine regulations on the dynamic progression of macrophages and could be a new promising therapy for endometriosis.

**Summary:** Niclosamide tunes the dynamic progression of peritoneal macrophages and their intercellular communications with B cells to rescue the disrupted microenvironment in the peritoneal cavity in a mouse model of endometriosis.

Graphic Abstract

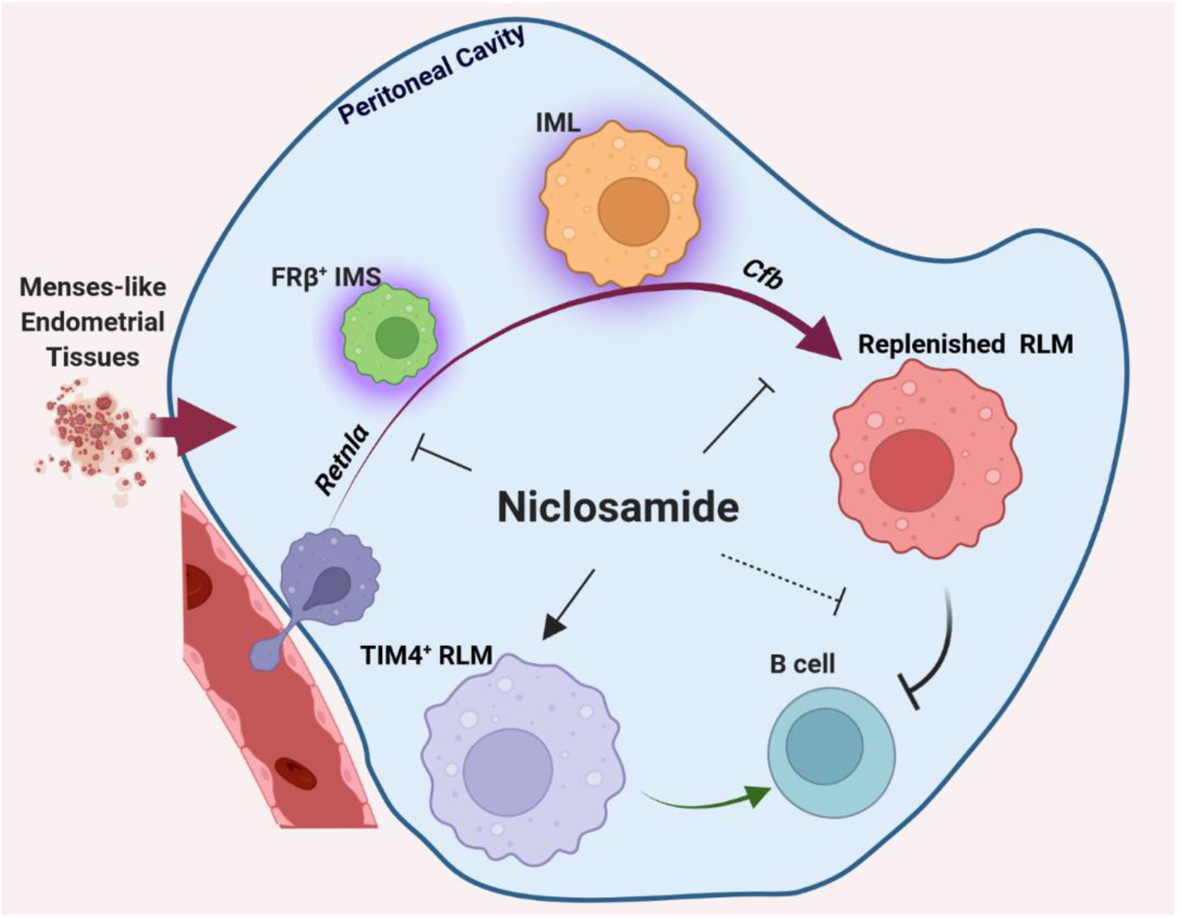

## Introduction

Endometriosis is a common chronic inflammatory disease that affects roughly 10% of reproductive-aged and adolescent women worldwide (Zondervan et al., 2018; Zondervan et al., 2020). It is characterized by the presence and growth of tissues resembling endometrium, termed endometriotic lesions, outside of the uterus. Patients with endometriosis exhibit symptoms of chronic pelvic pain, infertility, and multiple other health issues leading to tremendous reductions in their quality of life (Zondervan et al., 2020). Unfortunately, public and professional awareness of this disease remains poor. Current hormonal therapies, along with laparoscopic surgery, do not cure the disease and are often of limited efficacy with high recurrence rates, frequent side effects, and potential morbidity. Thus, a critical need exists to develop new and effective therapies for endometriosis targeting biologically important mechanisms that underlie the pathophysiology of this disease.

Disruption of the immune homeostasis in the peritoneal cavity drives the disease development of endometriosis, and macrophages play a central role in this process (Capobianco and Rovere Querini, 2013; Hogg et al., 2020; Shi et al., 2021). Peritoneal macrophages infiltrate endometriotic lesions and promote their growth and vascularization by releasing proinflammatory cytokines and growth factors (Bacci et al., 2009; Cheong et al., 2002; Sekiguchi et al., 2019; Shi et al., 2021). In addition, IGF1 and netrin-1, along with cytokines secreted by macrophages, also promote neurogenesis and innervation at lesion sites, which enhances the pain sensation in patients (Ding et al., 2021; Forster et al., 2019; Greaves et al., 2015; Scholl et al., 2009). The proinflammatory cytokines released by macrophages disrupted in endometriosis also affect multiple important activities of reproduction, such as hormonal balances and decidualization, leading to infertility (De Ziegler et al., 2010; Rasheed and Hamid, 2020). Suppressing the release of proinflammatory cytokines and growth factors from macrophages inhibits lesion growth and endometriosis­associated pain in rodent models (Bacci et al., 2009; Forster et al., 2019; Liu et al., 2018; Shi et al., 2021). Therefore, targeting peritoneal macrophages that are critical for maintaining immune homeostasis in the pelvic cavity could be a new direction for drug development in endometriosis therapy.

To fully characterize the role of peritoneal macrophages in the pathophysiology of endometriosis, a better understanding of the heterogeneity of macrophage populations and their subtype-specific contributions to endometriosis is necessary. Two subsets of macrophages have previously been characterized in the peritoneal cavity and are referred to as “small” (SPMs) and “large” peritoneal macrophages (LPMs) based on both their sizes and frequency (Ghosn et al., 2010). MHC II^high^ F4/80^low^ SPMs are short-lived and are recruited from Ly6C+ classical monocytes, while MHC II^low^ F4/80^high^ LPMs are resident and long-lived with an embryonic origin (Bain et al., 2016; Kim et al., 2016). The population of embryo-derived resident LPMs (RLMs) uniquely express TIM4, and its number is mainly maintained through its self-renewal under physiological conditions (Rosas et al., 2014). With mild inflammation, some of the recruited SPMs gradually differentiate into F4/80^high^ macrophages but still remain in an immature state due to the existence of RLMs (Bain et al., 2016). When the population of RLMs is ablated with extended inflammation, these transitory F4/80^high^ macrophages finally mature and replenish the pool of RLMs (Bain et al., 2016; Liu et al., 2019). However, this newly recruited resident population shows striking functional differences from those embryo-derived ones and thus increasing the risks for the incidence and severity of diseases in the future (Bain et al., 2016). In endometriosis, dynamic and progressive alterations of peritoneal macrophages were also found associated with lesion development (Hogg et al., 2021; Johan et al., 2019). However, the transcriptomic characteristics of peritoneal macrophages especially those transitory subtypes and the molecular signaling networks that coordinate the dynamic progression of macrophages in endometriosis are unknown.

Niclosamide is an FDA-approved anthelmintic drug with multiple clinical trials ongoing to repurpose it for the treatment of other diseases including cancer and metabolic diseases (Chen et al., 2018). We previously reported that niclosamide reduced lesion growth, alleviated aberrant inflammation in peritoneal fluids, and decreased the vascularization and innervation in lesions using mouse models of endometriosis (Prather et al., 2016; Shi et al., 2021).

In this study, we further focused on the heterogeneity of peritoneal macrophages and molecular mechanisms regulating their dynamic progression after lesion induction using a mouse model of endometriosis. Moreover, we found that niclosamide finely reversed those transcriptomic changes of macrophages caused by lesion induction through its stepwise regulations on the differentiation of recruited macrophages, the maturation of transitory LPMs, and the preservation of embryo-derived RLMs. Niclosamide also rescued the communications between LPMs and B cells which were disrupted by lesion induction. Therefore, we propose that macrophages could be the direct target of niclosamide, and niclosamide could be a new promising therapy for the treatment of endometriosis. Finally, to share our scRNA data with other researchers, we have created a cloud-based web tool (Webpage: https://kanakohayashilab.org/hayashi/en/mouse/peritoneal.immune.cells/) for the gene of interest searches that can be easily conducted without the requirement of complicated computer programming skills.

## Results

### Single-cell transcriptomic sequencing of peritoneal immune cells

In this study, endometriosis-like lesions (ELL) were induced by inoculating mense­like tissues from donor mice into the peritoneal cavity of the recipient, as described in the Method section. Three weeks later, one group of mice was administrated niclosamide (ELL_N) while the others (sham and ELL) were given a control vehicle (Fig. 1A). After another three weeks of treatment, cells in the peritoneal cavity, mostly immune cells, were collected and processed for single-cell transcriptomic analysis. A total of 13,679 cells with a median of 3,160 genes per cell were retained for downstream analysis after quality control and removal of low-quality cells.

**Figure 1.**
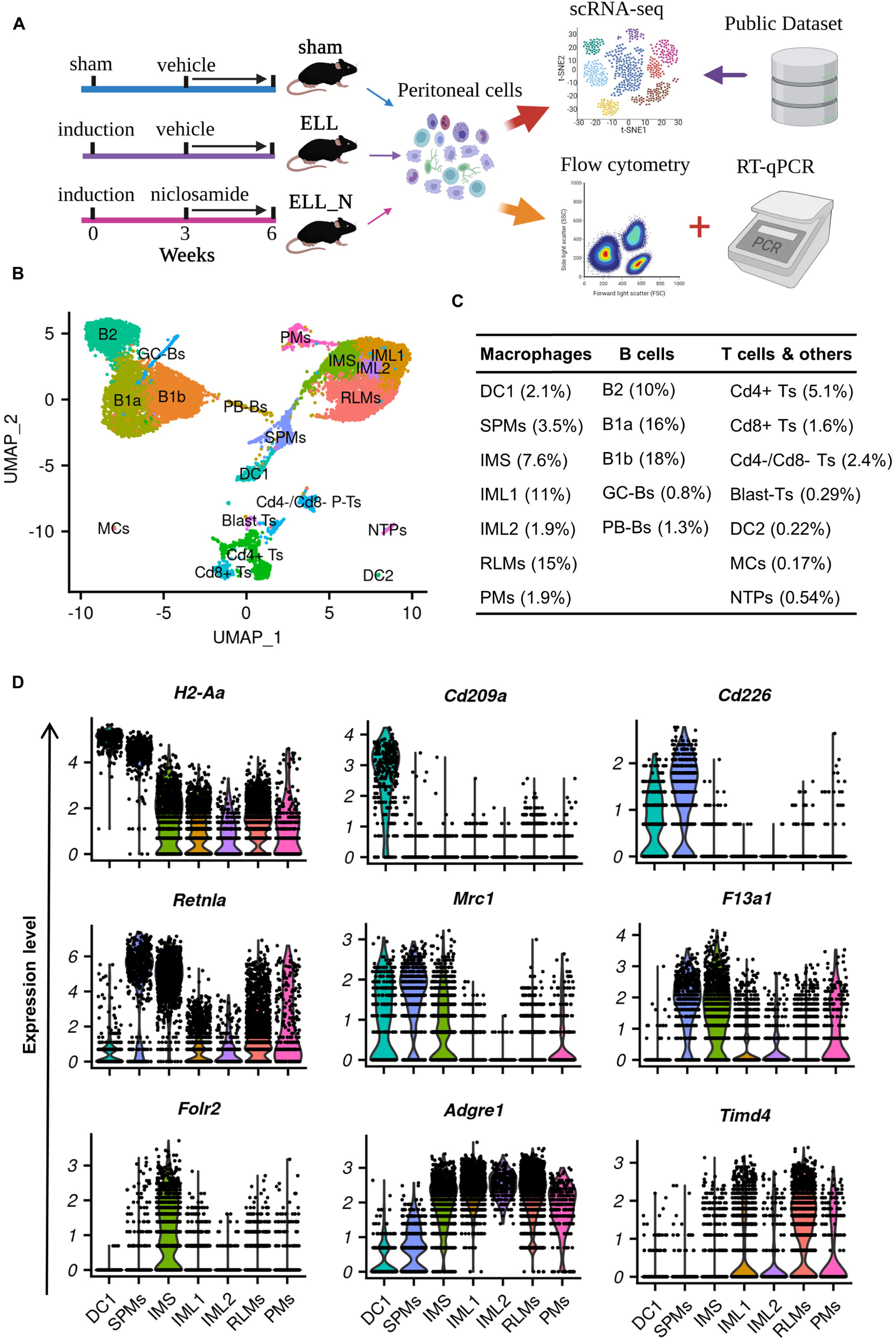
Single-cell transcriptomic profiling of peritoneal immune cells. (A) Schematic of the experimental pipeline. (B) UMAP visualization of peritoneal immune cells. (C) Ratios of each population in cell number. (D) VlnPlot of characteristic gene expressions for macrophage subpopulations. Sham, sham control; ELL, endometriosis-like lesions; ELL_N, niclosamide administration to ELL-induced mouse; DC1, dendritic cells 1; SPMs, “small” peritoneal macrophages; IMS, intermediate “small” macrophages; IML1, intermediate “large” macrophages subtype 1; IML2, intermediate “large” macrophages subtype 2; RLMs, resident “large” macrophages; PMs, proliferating macrophages; B1a, B1a cells; B1b, B1b cells; B2, B2 cells; GC-Bs, germinal-center B cells; PB-Bs, plasma blast B cells; Cd4+ Ts, Cd4+ T cells; Cd8+ Ts, Cd8+ T cells; Cd4-/Cd8-Ts, Cd4-/Cd8-T cells; Blast Ts, blast T cells; DC2, dendritic cells 2; MCs, Mast cells; NTPs, neutrophils.

Integrated cells from all three groups of samples were classified into 19 clusters based on the unsupervised clustering workflow of the Seurat package with cell identities determined by canonical marker gene distributions (Fig. 1B and S1A). Cells from each group showed a consistent distribution of each cluster in UMAP (Fig. S1B), suggesting an unbiased capture of cell populations between groups of different treatments. Macrophages (43%) and B cells (46%) are the most abundant populations identified in peritoneal fluids along with much fewer T cells and other immune cells (Fig. 1B and C).

### Heterogeneity of peritoneal macrophage populations

Interestingly, 7 sub-clusters of macrophage-related populations (DC1, SPMs, IMS, IML1, IML2, RLMs, and PMs) were identified in these samples (Fig. 1B and C). Clusters of dendritic cells 1 (DC1) and “small” peritoneal macrophages (SPMs) express high levels of characteristic SPMs markers including *H2-Aa* and *Irf4* and are distinguished by the expression of *Cd209a* in DC1 and *Cd226* in SPMs (Fig. 1D and S1C). The identity of resident “large” peritoneal macrophages (RLMs) was determined by the expression of its unique marker, *Timd4* (encodes TIM4, Fig. 1D). In addition, a group of proliferating macrophages (PMs) was identified by their exclusive expression of proliferating markers, *Mki67* and *Birc5* (Fig. S1C).

In addition to these well-known types of peritoneal macrophages, three novel subtypes: intermediate “small” macrophages (IMS), intermediate “large” macrophages 1 (IML1), and intermediate “large” macrophages 2 (IML2), were also identified. Cells of IMS showed unique expression of *Folr2* (encodes folate receptor β (FRβ) subunit, Fig. 1D). Though cells of IMS showed expression of LPM markers including *Adgre1* (F4/80) and *Icam2* (CD102), equivalent levels of *Retnla* (also known as *Relmα* or *Fizz1*), *Mrc1* (CD206), *F13a1*, and *Aif1* as cells of the SPMs were also found in the IMS (Fig. 1D and S1C). These gene expression characteristics of IMS suggest their close developmental relations to SPMs. The other two intermediate groups (IML1 and IML2) showed high expression of *Adgre1* and *Icam2*, but they are low in the expression of *Timd4*, indicating that they are newly recruited immature LPMs.

Next, the top 100 differentially expressed genes in each cluster were used to enrich their unique characteristics by gene ontology analysis (GO) of biological processes (Fig. S2 and Table S1). Up-regulated biological processes related to TNF production were found in all three intermediate subtypes (IMS, IML1, and IML2) based on enriched terms of “regulation of tumor necrosis factor production” in IMS, “positive regulation of tumor necrosis factor production” and “negative regulation of transforming growth factor beta production” in IML1, and “tumor necrosis factor production” in IML2. Different from these intermediate subtypes, one term of “regulation of transforming growth factor beta production” was enriched in RLMs, suggesting differential functions between monocyte-derived and embryo-derived macrophages in endometriosis. In addition, the three intermediate groups all show characteristics of “phagocytosis” or “cell killing”, which were not found in the population of RLMs. Different from DC1 and SPMs, enriched biological processes to support the “regulation of angiogenesis” were found in the cells of IMS, IML1, IML2, and RLMs. This high resolution analysis of macrophage transcriptomes identified in our study provides us with an unprecedented opportunity to study stage-specific effects caused by ELL and the treatment of niclosamide.

### Peritoneal macrophages in a normal physiological state

A publicly-available single-cell RNA-seq dataset from CD11b+ peritoneal macrophages in wild-type female mice in a normal physiological state [GSM4151331, (Bain et al., 2020)] was re-analyzed in this study (Fig. S3A and B). Similar subpopulations related to the “small” peritoneal macrophage lineage were identified and named “DC”, “SPM”, and “IMS”. Four clusters of *Timd4*+ embryo-derived resident macrophages were also identified (RLM1-4). However, different from our samples collected from Sham, ELL, and ELL_N groups, no intermediate “large” (IML) subtypes were distinguished in these macrophages at physiological state. However, their results also support that the continuous differentiation of recruitment macrophages was driven by disrupted homeostasis of macrophages under external stimuli like lesion induction.

### Niclosamide reverses lesion-induced transcriptomic changes in macrophages

Transcriptomic changes in the macrophages induced by ELL and niclosamide (ELL_N) were compared by the gene set enrichment analysis (GSEA). Compared to the sham group, a total of 135 biological processes were up-regulated by ELL (Fig. 2A and Table S2), and 78 of them were reversed by niclosamide (Fig. 2A and Table S2). More specific, ELL induced activation, proliferation, and differentiation of peritoneal macrophages as GO terms of “Establishment or maintenance of cell polarity”, “Transmembrane receptor protein tyrosine kinase signaling pathway”, “Lymphocyte proliferation”, “Positive regulation of cell migration” and “Regulation of lymphocyte differentiation” were all positively enriched compared to the sham group (Fig. 2D). ELL also up-regulated biological processes of “Vesical organization”, “Ceramide transport”, “Regulation of neuron differentiation”, and “Angiogenesis”, which are associated with vascularization, neurogenesis, and pain sensation in lesions. All of the biological processes above were suppressed by niclosamide compared to the ELL group (Fig. 2D). On the other side, 10 out of 12 ELL-inhibited pathways were enhanced by niclosamide (Fig. 2B and Table S2). For example, niclosamide promoted biological processes of “Cytoplasmic translation” and “Oxidative phosphorylation” which were reduced by ELL (Fig. 2E and Table S2).

**Figure 2.**
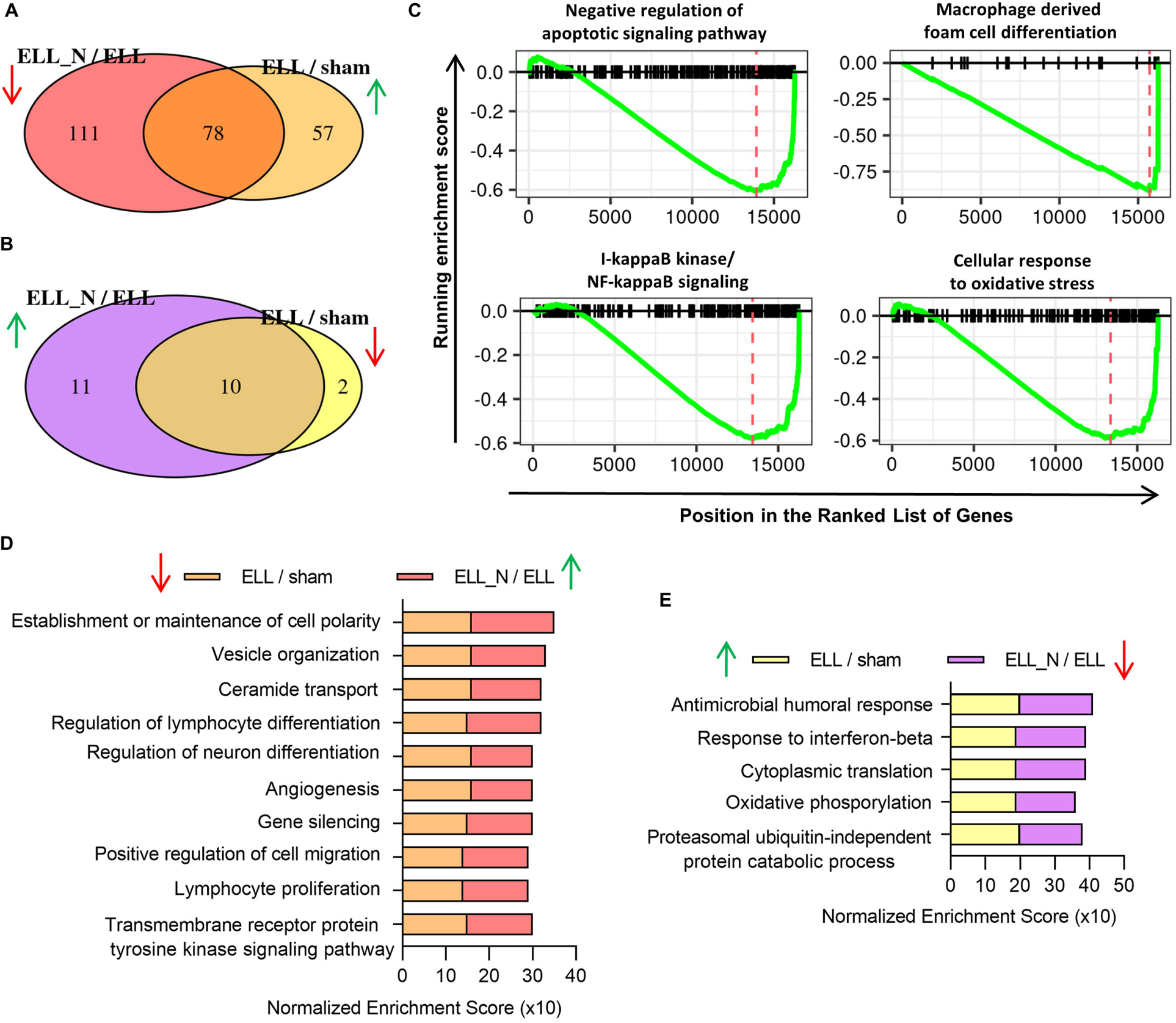
Niclosamide finely reverses transcriptomic changes caused by endometriosis-like lesions (ELL) to peritoneal macrophages. (A) Venn diagram shows the overlaps of enriched GSEA terms of biological processes between those enhanced in by ELL (ELL/sham) and those reduced by niclosamide (ELL_N/ELL). (B) Similar to (A) but shows the overlaps between enriched terms of those down-regulated in ELL (ELL/sham) and up-regulated by niclosamide (ELL_N/ELL). (C) Enriched GSEA terms of biological processes that were reduced within the ELL_N group compared to the sham group. (D) Representative GSEA terms that were enhanced by ELL (ELL/sham) and were decreased by niclosamide (ELL_N/ELL). (E) Representative GSEA terms that were inhibited by ELL (ELL/sham) and were reversed by niclosamide (ELL_N/ELL).

Niclosamide tunes disrupted macrophages back to a relevant homeostatic level with the sham group after 3 weeks of treatment (ELL_N/sham), with only 25 differential regulated pathways found (Fig. 2C and Table S2). Compared to the sham group, niclosamide further decreased the inflammatory responses and oxidative stress in macrophages but promoted their apoptosis as indicated by enriched GO terms of “Macrophage derived foam cell differentiation”, “I-kappaB kinase/NF-kappaB signaling”, “Cellular response to oxidative stress” and “Negative regulation of apoptotic signaling pathways” (Fig. 2C). These results indicate that ELL induced aberrant activation of macrophages and enhanced their signaling communications for lesion growth and pain sensation, which were finely reversed by niclosamide at the transcriptomic level.

### Niclosamide suppressed the expression of genes that were enhanced by ELL

To further understand the transcriptomic changes in macrophages caused by ELL and niclosamide, we examined their corresponding alterations at the gene level (Fig. 3A). A total of 116 genes were up-regulated by ELL compared to the sham group (ELL/sham) with 91 of them being decreased by niclosamide (ELL_N/ELL). These top up-regulated genes by the presence of ELL include *Retnla*, *Mrc1*, *F13a1*, *Kctd12*, *Plxnd1*, *Ccl9*, *Ccl6*, and *Socs6*, which were all subsequently inhibited by niclosamide (Fig. 3C). These differentially expressed genes were also confirmed by qPCR analyses in independent samples (Fig. 3D). Consistently, ELL enhanced the expression of *Retnla*, *Mrc1*, *Ccl6*, *Kctd12,* and *Scocs6* while niclosamide suppressed the expression of *Retnla*, *F13a1*, *Mrc1*, *Ccl6 (p=0.06)*, *Kctd12*, *Scocs6*, and *Plxnd1.* Interestingly, most of these up-regulated genes by ELL were uniquely distributed in the populations of DC1, SPMs, and IMS (Fig. 3E) but not the LPMs subtypes (IML1, IML2, RLMs). Consistent with their distributions in our samples, a similar pattern was also found in the re-analyzed dataset of macrophages at the normal physiological state (DC, SPM, and IMS; Fig. S3C). Therefore, ELL enhanced gene expressions related to the lineage of recruited “small” macrophages, and niclosamide reversed those changes.

**Figure 3.**
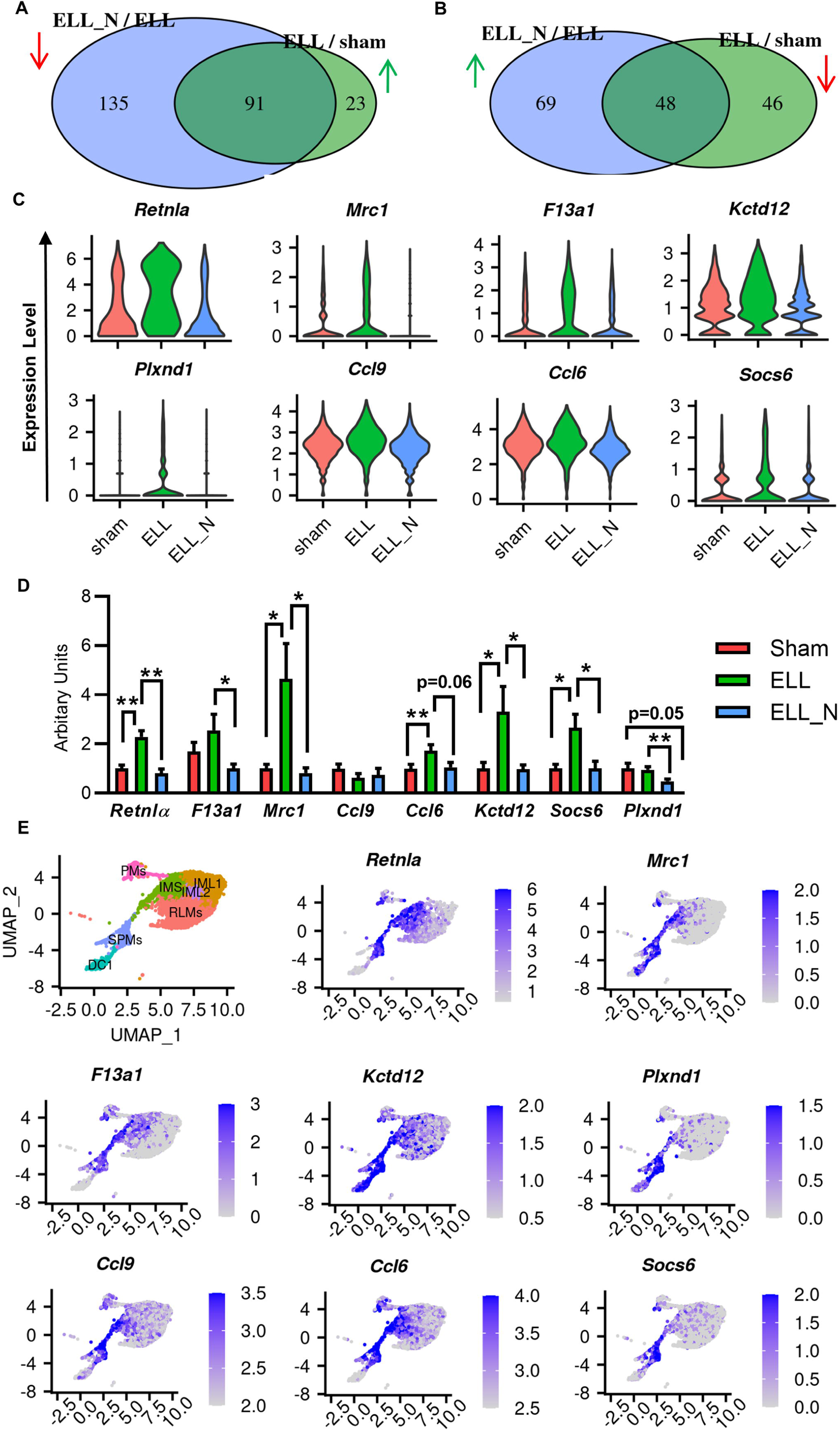
Niclosamide up-regulated genes with specific distributions within the “small” peritoneal macrophage lineage. (A) Venn diagram shows that overlaps of genes that were enhanced by ELL (ELL/sham) but decreased by niclosamide (ELL_N/ELL). (B) Overlaps of genes that were inhibited by ELL (ELL/sham) but enhanced by niclosamide (ELL_N/ELL). (C) Vlnplot shows representative genes that were enhanced by ELL but were reduced by niclosamide treatment (ELL_N). (D) Verification of differential gene expression by RT-qPCR. \**p* < 0.05, \*\**p* < 0.01,\*\*\**p* < 0.001, mean ± SEM, n = 6 per each group. (E) UMAP of computationally selected macrophage populations and the distribution of genes within them.

### Niclosamide attenuates the recruitment of “small” peritoneal macrophages

As cells from DC1, SPMs, and IMS showed close developmental relations based on their gene expression patterns (Fig. 1D and S1C), the differentiation of cells from DC1 to SPMs and IMS is considered a continuous process. Therefore, we next applied pseudo-temporal trajectory analysis to these three subpopulations to elucidate the dynamic changes and functions of ELL-enhanced genes along this early differentiation process of recruited macrophages. Cells from DC1, SPMs, and IMS were computationally selected, and a continuous trajectory of the differentiation process was constructed using Monocle3 (Fig 4A). The expression of genes enhanced by ELL including *Retnla*, *Mrc1*, *F13a1*, *Kctd12*, *Plxnd1*, *Ccl9*, *Ccl6,* and *Socs6* (Fig. 3B and C) were plotted along this developmental timeline (Fig. 4B). Interestingly, most of these genes showed a consistent upregulation pattern during the early differentiation process of SPMs from DC1 (Fig. 4B). This dynamic expression pattern was also confirmed by the re-analyzed datasets of macrophages at physiological states (Fig. S4A and B).

**Figure 4.**
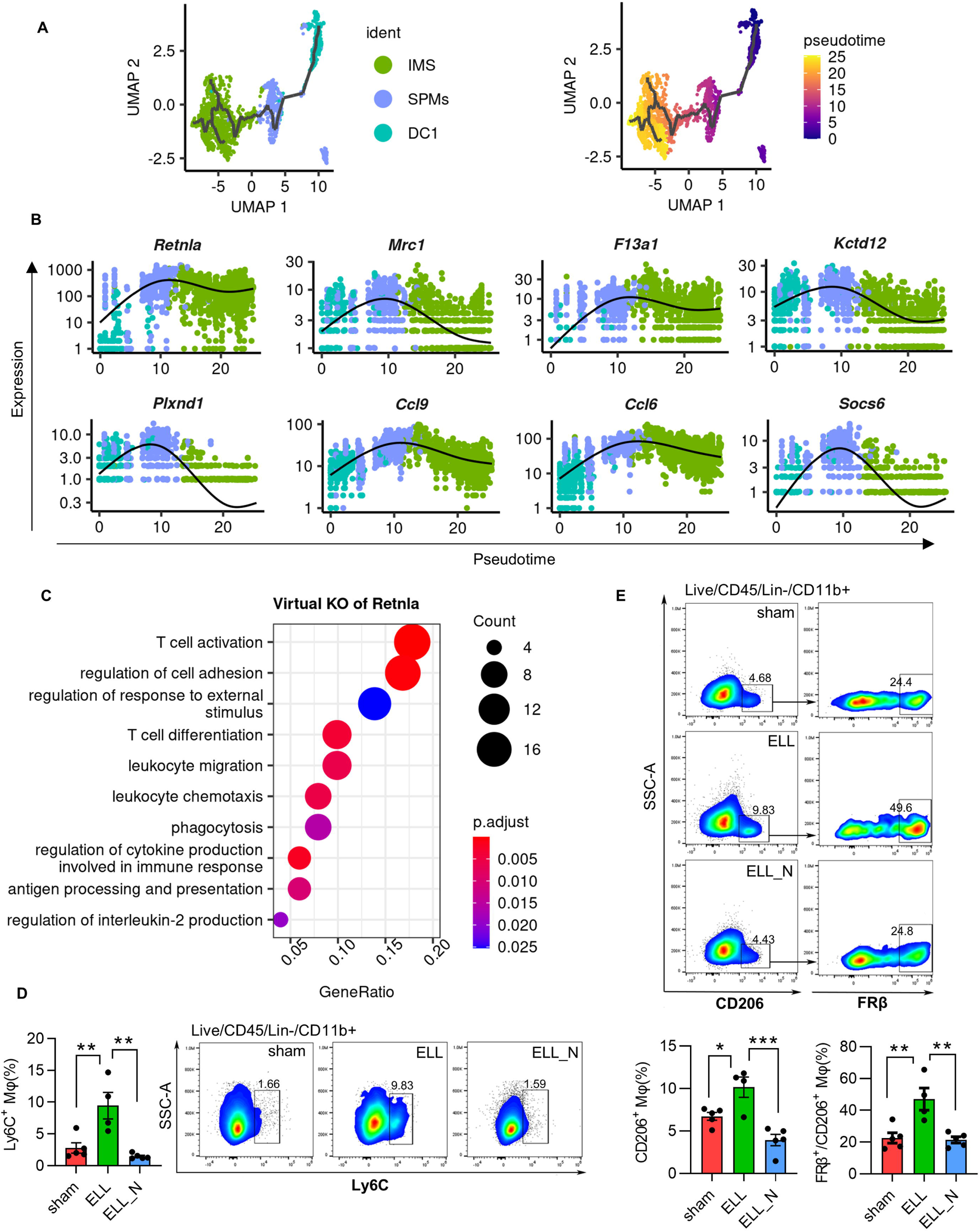
Reconstruction of a pseudo-temporal trajectory for the early differentiation of recruited macrophages. (A) UMAP shows selected cells of recruited “small” macrophages (left) and a trajectory path built by Monocle 3 (right). (B) Dynamic changes of genes along this trajectory path of differentiation. (C) GO terms of biological processes that were enriched by perturbed genes affected by virtual KO of *Rentla*. (D) Flow cytometer isolation and quantification of Ly6c+ recruited monocytes. (E) Flow cytometer results and quantification of CD206+ and FRβ+ macrophages. \**p* < 0.05, \*\**p* < 0.01, mean ± SEM, n = 5 per each group.

Among those genes, the expression of *Retnla* increases by about 100-folds during this process of differentiation (Fig. 4B). The function of *Retnla* for this dynamic process was further studied by *in silico* knockout *Retnla* in these three subpopulations. The top dysregulated genes by virtual knockout of *Retnla* include *Gas6*, *H2-Oa*, *Cd209a*, *Ccr2*, *Il1b,* and *Folr2* (Table S3). Those perturbed genes affected multiple important pathways of immune responses such as biological processes of “leukocyte migration”, “leukocyte chemotaxis”, “phagocytosis”, “regulation of cytokine production involved in immune response”, “antigen processing and presentation” and “regulation of interleukin-2 production” (Fig. 4C and Table S3). In addition, *Retnla* knockout may also affect the communications between macrophages and T cells, shown by GO terms of “T cell activation” and “T cell differentiation”.

As ELL enhanced the expression of genes that are necessary for the differentiation of SPMs from DC1, an increased population number of SPMs was expected in the ELL group. By the analysis of flow cytometry, we confirmed a more than 3 times increase in the number of Ly6C+ recruited small macrophages in the group of ELL, which was reduced to a similar level with the sham group by niclosamide (Fig. 4D). As a consequence, the populations of FRβ+ (encoded by *Folr2*) IMS and CD206+ (encoded by *Mrc1*) recruited macrophages were also increased by ELL and decreased by niclosamide (Fig. 4E). Therefore, niclosamide attenuated the recruitment of macrophages by decreasing its differentiation from DC1, which was promoted by ELL.

### Niclosamide increased the expression of genes that were decreased by ELL

In addition to niclosamide’s suppressions on the genes that were enhanced by ELL, niclosamide also up-regulated over 50% of genes that were reduced by ELL induction (Fig. 3B). The top representative genes include *Cfb*, *Hp*, *Ifitm2*, *Ifitm3*, *Gbp2b*, *C1qb*, *Prdx5,* and *Gngt2* (Fig. 5A). The results of qPCR of immune cells in the peritoneal fluid with different treatments showed consistent changes in the expression of *Cfb*, *Ifitm2*, *Ifitm3*, *Gbp2b*, *C1qb*, *Prdx5,* and *Gngt2* (Fig. 5B). Interestingly, most of these genes were highly expressed in the intermediate subtypes of macrophages and RLMs, but their expression in DC1 and SPMs was very low (Fig. 5C).

**Figure 5.**
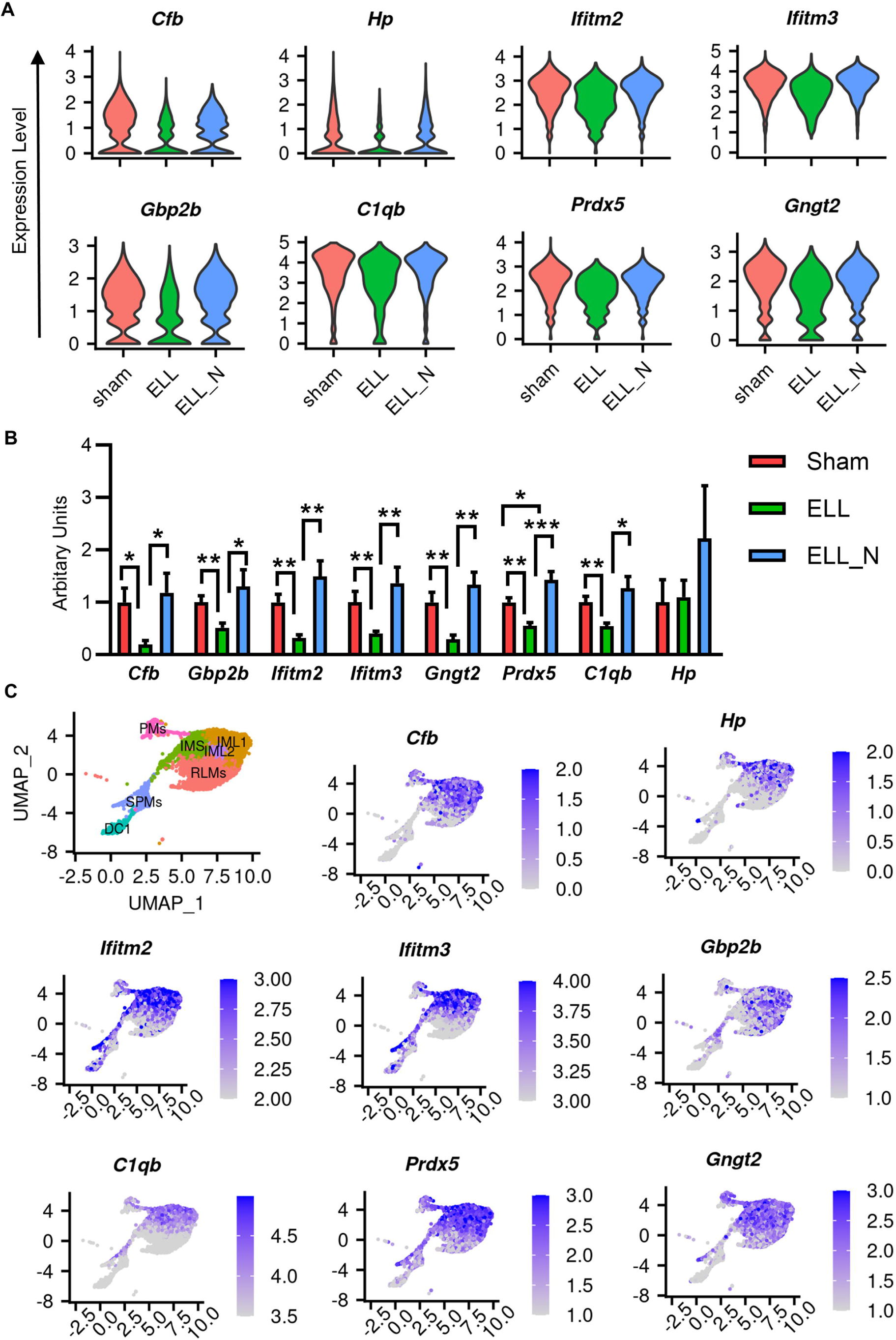
Niclosamide downregulated genes with specific distributions within the “large” peritoneal macrophage lineage. (A) Vlnplot shows representative genes that were reduced by ELL (ELL/sham) but were enhanced by niclosamide (ELL_N/ELL). (B) Verification of differential gene expression by RT-qPCR. \**p* < 0.05, \*\**p* < 0.01, \*\*\**p* < 0.001, mean ± SEM, n = 6 per each group. (C) UMAP of computationally selected macrophage populations and the distribution of genes within them.

### Niclosamide decreased the maturation of recruited macrophages

Inflammatory conditions lead to the recruitment of LPMs from bone marrow which gradually mature and replenish the resident macrophage pool though they would still be functionally different from those embryo-derived resident macrophages (Louwe et al., 2021). The three subtypes of “large” peritoneal macrophages, IML1, IML2, and RLMs (Fig. 6A) were involved in this biological process of maturation and replenishment and were used to reconstruct a pseudo developmental trajectory (Fig. 6A). Then, the down-regulated genes caused by ELL including *Cfb*, *Hp*, *Ifitm2*, *Ifitm3*, *Gbp2b*, *C1qb*, *Prdx5,* and *Gngt2* (Fig. 5A) were plotted along this trajectory path (Fig. 6B). Interestingly, the expression of these genes showed a consistent decreasing pattern during this biological process, suggesting their important roles in maturation inducing.

**Figure 6.**
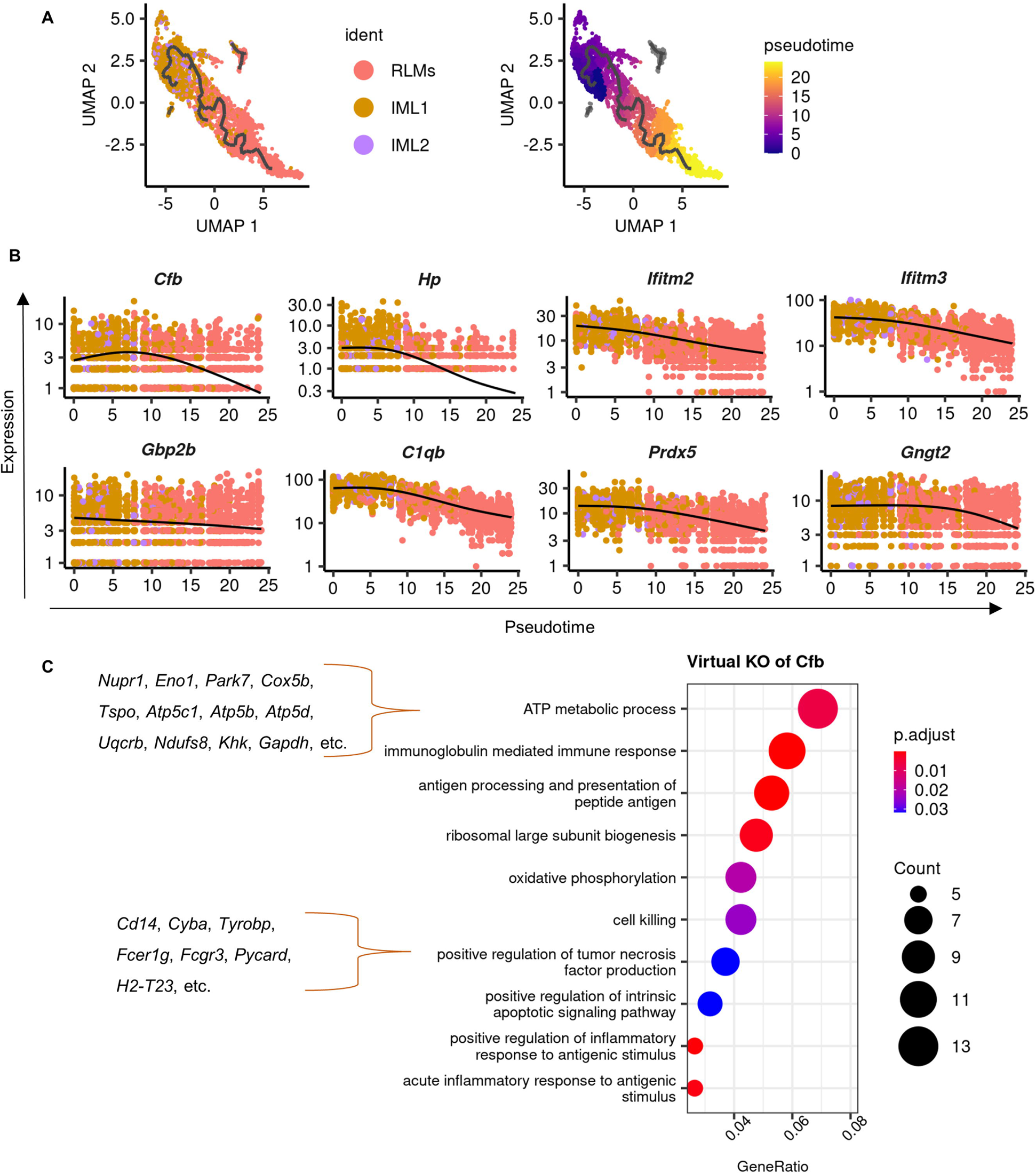
Reconstruction of a pseudo-temporal trajectory for maturation of intermediate “large” macrophages. (A). UMAP shows selected cells of “large” macrophages (left) and a trajectory path built by Monocle 3 (right). (B). Dynamic changes of genes along this trajectory path of maturation and replenishment. (C). GO terms of biological processes that were enriched by perturbed genes affected by virtual KO of *Cfb*.

As *Cfb* is one of the most responsive genes regulated by ELL and niclosamide in this biological process (Fig. 5A), we further explored its functions by *in silico* knockout of *Cfb* in cells of IML1, IML2, and RLMs (Table S3). This analysis showed that virtual knockout of *Cfb* not only disrupted the inflammatory responses of macrophages but also changed the metabolism, protein synthesis, and apoptosis of macrophages as terms of “ATP metabolic process”, “Oxidative phosphorylation”, “ribosomal large subunit biogenesis”, and “positive regulation of intrinsic apoptotic signaling pathway” were enriched based on genes disrupted by *Cfb* knockout (Fig. 6C and Table S3). Moreover, *Cfb* seems to be important for TNF production in these LPMs based on enriched terms of “positive regulation on tumor necrosis factor production” (Fig. S2).

As no intermediate “large” macrophage phenotypes were identified in the public-available dataset of macrophages at physiological states, no obvious changes of genes mentioned above were found along the trajectory built between those LPMs (LPM1-4, Fig. S5A). This difference also supports the previous finding that the maturation of macrophages is only active upon external stimuli. Therefore, ELL promotes the maturation of recruited macrophages to replenish the resident macrophage pool by downregulating genes such as *Cfb*, and this process is inhibited by niclosamide.

### Niclosamide reduces the ablation of embryo-derived resident macrophages

The existence of embryo-derived RLMs prohibits the replenishment of the resident macrophage pool by recruited ones, while increased inflammation leads to ablation of embryo-derived RLMs and promotes the process of replenishment (Louwe et al., 2021). In support of this, ELL decreased the expression of *Timd4* and *Apoc1*, markers for embryo-derived RLMs, in the peritoneal macrophages (Fig. 7A and B). Expression of *Timd4* and *Apoc1* was also found to increase along with the initial maturation process of intermediate macrophages, suggesting recruited macrophages also gradually acquire their residency (Fig. 7C). Furthermore, knockout of *Timd4* in these LPM populations was shown to induce their B cell characteristics as terms of “B cell activation”, “B cell receptor signaling pathway” and “B cell proliferation” were enriched based on disrupted genes of *Timd4* knockout (Fig. 7D and Table S3). Knockout *Timd4* also induced changes in TNF production, phagocytosis, and oxidative stress in those “large” types of macrophages.

**Figure 7.**
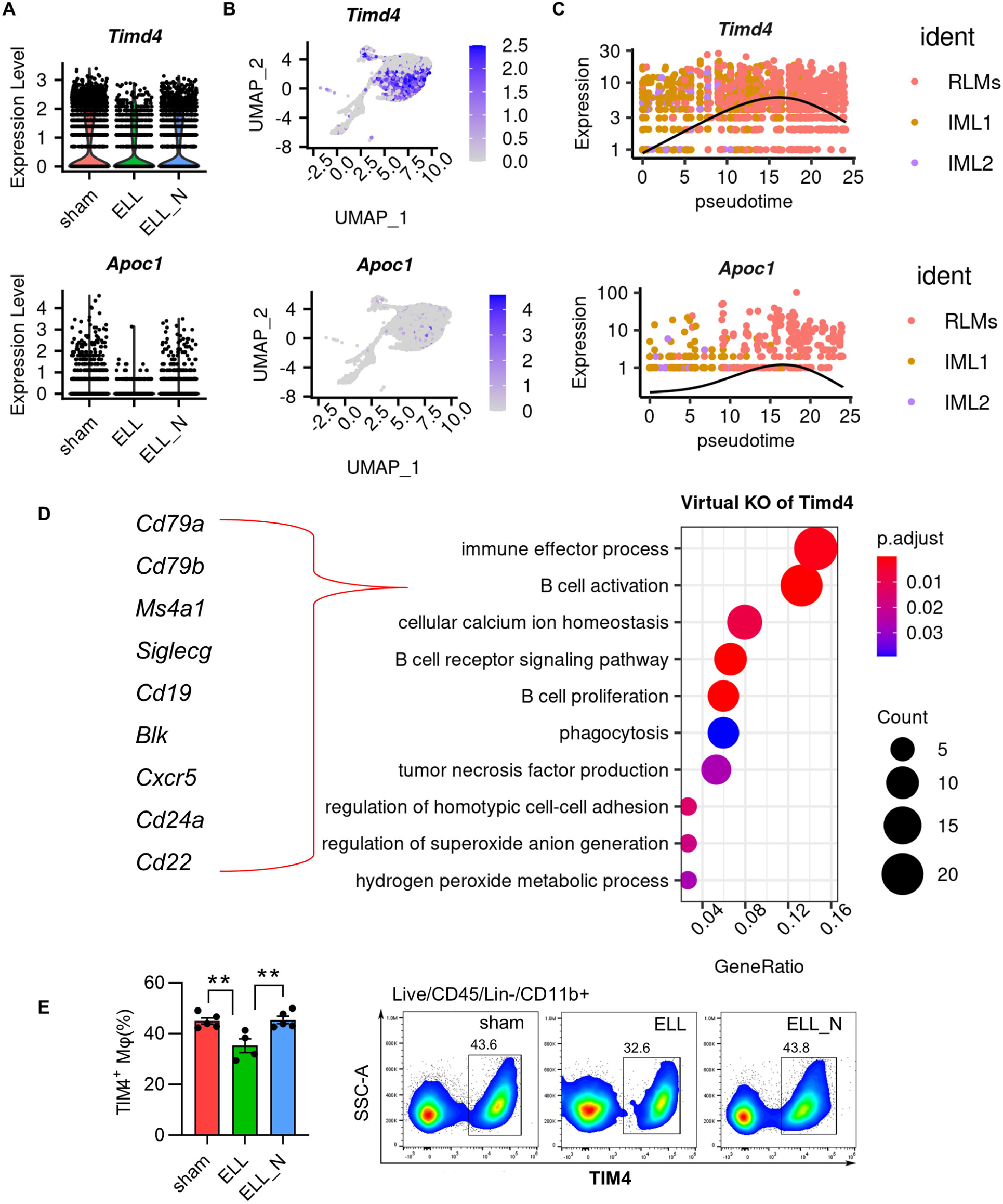
Niclosamide preserves the population of embryo-derived resident “large” peritoneal macrophages (RLMs). (A) Vlnplot shows gene expression of *Timd4* and *Apoc1*. (B) Feature plot shows the distribution of *Timd4* and *Apoc1*. (C) Dynamic changes of genes along the trajectory path of maturation and replenishment (D) GO terms of biological processes that were enriched by perturbed genes affected by virtual KO of *Timd4*. (E) Flow cytometer isolation and quantification of TIM4+ resident macrophages.\*\**p* < 0.01, mean ± SEM, n = 5 per each group.

As a consequence of reduced *Timd4* expression in macrophages by ELL, the population number of TIM4+ RLMs was also reduced (Fig. 7E). Niclosamide rescued the expression of *Timd4* and also the number of TIM4+ RLMs (Fig. 7E). Thus, ELL enhanced the replenishment of the resident macrophage pool by increasing the ablation of embryo-derived RLMs, which further promotes the maturation of recruited macrophages. Niclosamide preserves these embryo-derived RLMs by enhancing *Timd4* expression and recuses the homeostasis of macrophages disrupted by ELL.

### Niclosamide rescues the communications between macrophages and B cells

CXCL13-producing embryo-derived RLMs play an important role in the maintenance and recruitment of peritoneal B1 cells upon inflammation, while newly recruited macrophages were reported to be deficient in CXCL13 (Bain et al., 2020; Beattie et al., 2016; Louwe et al., 2021; Zeng et al., 2018). To understand the communications between macrophages and B cells among their subpopulations, two important signaling networks for immune cell recruitment, CXCL and CCL, were analyzed based on the ligand and receptor interactions between cells using the CellChat package. Most of the CCL ligands were found to be released by macrophages and received by themselves, whereas there were very few communications between macrophages and B cells (Fig. 8A). The SPMs were the most affected cells by the CCL signaling from the other types of macrophages, which is consistent with their increased recruitment by ELL induction.

**Figure 8.**
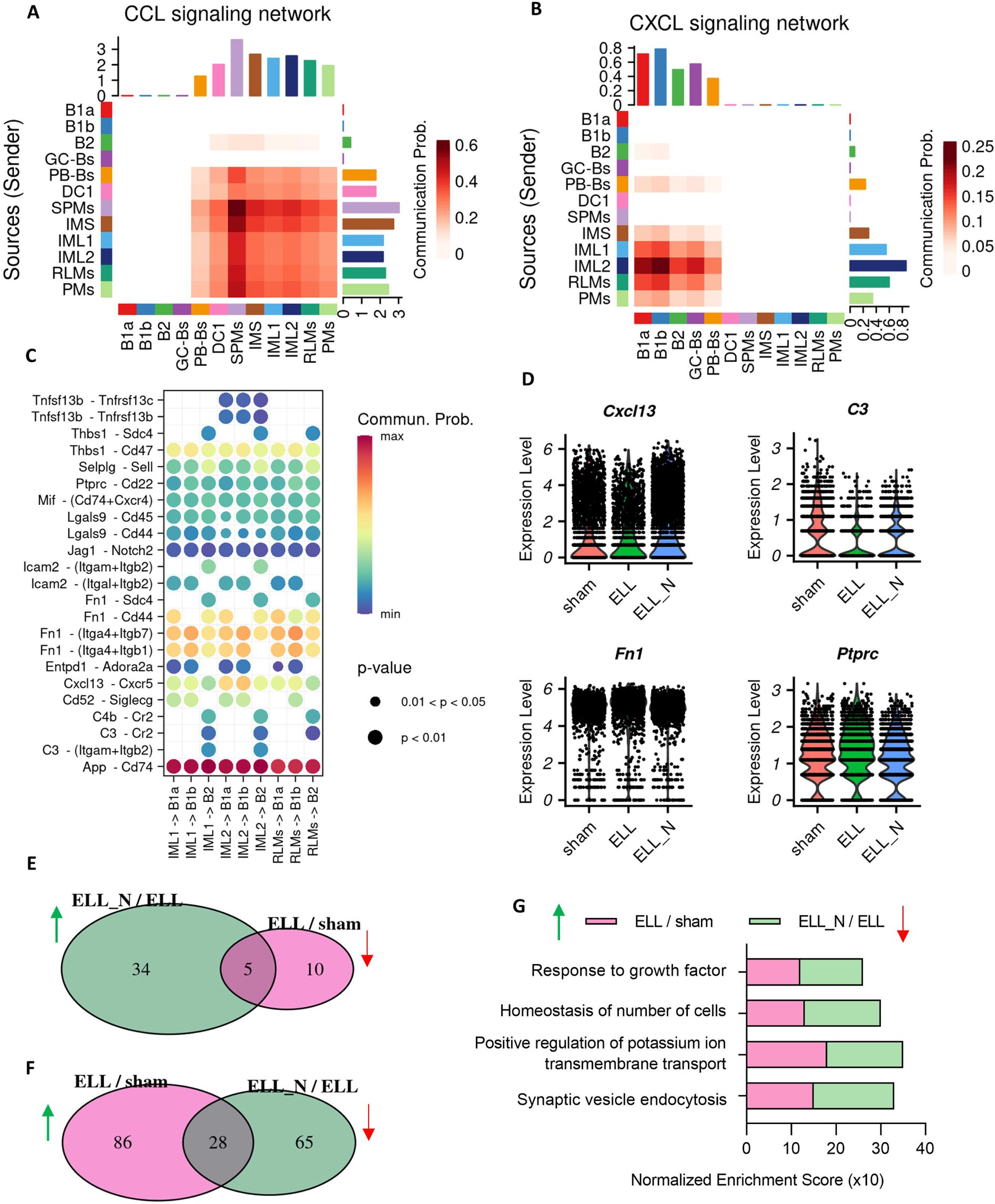
Niclosamide rescued the disrupted intercellular communications from macrophages to B cells. (A) Heatmap showing the interactions between macrophages and B cells in terms of CCL signaling networks. (B) Heatmap showing the interactions between macrophages and B cells in terms of CXCL signaling networks. (C) Bubble plot showing all significant expressed ligand-receptor pairs identified between three “large” macrophages (IML1, IML2, RLMs) and three subtypes of B cells (B1a, B1b, B2). (D) Vlnplot showing the differential expressed genes in macrophages. (E) Overlaps of GO biological processes in B cells that were suppressed by ELL (ELL/sham) but enhanced by niclosamide (ELL_N/ELL). (F) Overlaps of GO biological processes in B cells that were promoted by ELL (ELL/sham) but suppressed by niclosamide (ELL_N/ELL). (G) Overlaps of GO biological processes that were promoted by ELL by suppressed by niclosamide.

On the other hand, the expression of CXCL ligands was found to be mostly expressed in “large” types of macrophages, such as IML1, IML2, and RLMs, and signals received by B cells (Fig. 8B). Furthermore, we explored the interactions between these CXCL-producing “large” types of macrophages and the three largest populations of B cells (B1a, B1b, and B2). Twenty-three significantly expressed ligand-receptor pairs were identified including *App*-*Cd74*, *Cxcl13*-*Cxcr5*, *C3*-*Cr2*, *Fn1*-*Sdc4*, *Ptprc*-*Cd22*, which may play important roles in the recruitment and functionality of B cells under the inflammatory condition induced by ELL induction. Among them, decreased expression of *Cxcl13* and *C3* was found in the peritoneal macrophages by ELL, and their expression was enhanced by niclosamide (Fig. 8D). Moreover, B1a and B1b cells received the most *Cxcl13* signals, while B2 cells exclusively received the signals of *C3* (Fig. 8C). In addition, the expression of *Fn1* and *Ptprc* in macrophages was also altered by ELL and niclosamide (p<0.05), but the changes are limited.

The transcriptomic changes caused by ELL and niclosamide to B cells were also analyzed. Compared to the macrophages, the overlaps of biological processes that were disrupted by ELL (ELL/sham) and reversed by niclosamide (ELL_N/ELL) were much fewer (Figs. 8E and F; Table S4). Only 28 out of 114 biological processes that were enhanced by ELL were reversed by niclosamide (Fig. 8F; Table S4). The limited overlaps include biological processes of “response to growth factor” and “homeostasis of number of cells” (Fig. 8G), which may be associated with the recruitment and maintenance of the number of B cells through macrophages.

These results suggest that ELL disrupted the communications between “large” types of macrophages and B1/B2 cells, which were rescued by niclosamide through up-regulating the expression of *Cxcl13* and *C3*. The transcriptomic alternations caused by niclosamide to B cells are minimal, indicating that macrophages are possibly the direct targets of niclosamide.

## Discussion

Niclosamide is an FDA-approved oral anthelmintic drug that is originally used to treat human tapeworm infections (Selection et al., 2014). In addition to this common use, many clinical studies are ongoing to repurpose niclosamide for the treatment of other diseases such as different types of cancer, bacterial and viral infections, neuropathic pain, systemic sclerosis, and metabolic diseases (Chen et al., 2018; Fonseca et al., 2012; Jurgeit et al., 2012; Morin et al., 2016; Osada et al., 2011; Tao et al., 2014; You et al., 2014; Zhang et al., 2013). Though the clear direct binding targets of niclosamide have not been identified, studies have shown that niclosamide affects multiple important signaling pathways. One of the most appreciated action mechanisms is that niclosamide acts as a protonophore and thus uncouples oxidative phosphorylation and affects pH balance in cells (Chen et al., 2018; Jurgeit et al., 2012; Tao et al., 2014). In addition, signaling pathways of mTOR, Wnt/β-catenin, STAT3, NF-κB, and Notch are also modulated by niclosamide (Fonseca et al., 2012; Jin et al., 2010; King et al., 2015; Wang et al., 2009; You et al., 2014; Zhang et al., 2013). In endometriosis, we also found that niclosamide reduced the growth of lesions and decreased inflammation at lesions through its suppression of STAT3 and NF-κB signaling (Prather et al., 2016; Sekulovski et al., 2019; Sekulovski et al., 2020). Moreover, we also found that niclosamide does not disrupt reproductive functions in mice, making it a relatively safe drug for treatment (Prather et al., 2016).

Abnormal activation and increased numbers of macrophages along with elevated levels of proinflammatory cytokines such as IL-1β, IL6, IL8, and TNFα were found in the peritoneal fluid of patients with endometriosis (Milewski et al., 2008; Scholl et al., 2009; Zhou et al., 2020). We also reported that niclosamide treatment reduced inflammation in the peritoneal fluid as well as lesions, pelvic organs (uterus and vagina), and dorsal root ganglion (Shi et al., 2021). Consequently, niclosamide also decreased macrophage infiltration, vascularization, and innervation in the endometriotic lesions (Shi et al., 2021). In this study, we further reported that most of the transcriptomic changes in peritoneal macrophages induced by ELL were rescued by niclosamide. These changes include necessary biological processes for macrophage activation and for its communications with other types of cells, such as processes to promote angiogenesis and neurogenesis. Consistent with the unique function of niclosamide for mitochondria uncoupling, the biological processes of oxidative phosphorylation in macrophages were promoted by niclosamide. Furthermore, niclosamide rescues disrupted macrophage subpopulations back to a similar transcriptomic level compared with those in the sham group and further reduced the signaling of NF-κB and oxidative stress, and promoted apoptosis in macrophages after 3 weeks of treatment.

Previous studies have suggested the long-term existence of transitory macrophages in the peritoneal cavity under inflammation in addition to traditionally recognized SPMs and LPMs, but our knowledge of those immature subtypes and their contributions to disease development is limited (Bain et al., 2020; Liu et al., 2019; Louwe et al., 2021). In this study, we further characterized these subpopulations and identified three intermediate immature subtypes, named IMS, IML1, and IML2. Cells of the IMS showed close developmental relationships with SPMs with high expression of *Retnla* (*Relmα* or *Fizz1*), *Mrc1* (CD206), *F13a1*, and *Aif1*, which are important markers for monocyte-derived cells (Elizondo et al., 2019; Lee et al., 2014; Porrello et al., 2018; Yu et al., 2020). The populations of IML1 and IML2 are quite similar in transcriptomes and are F4/80^high^ MHC II^low^, but both of them are low in the expression of *Timd4*, suggesting that they are newly recruited LPMs. All three intermediate groups were characterized by TNF production, while the embryo-derived RLMs are supportive of TGFβ production. In addition to TNF production, cells of IMS also support angiogenesis. Therefore, their prolonged existence by ELL may promote lesion growth.

Under a successful resolution of inflammation, the recruited SPMs decrease their numbers by apoptosis or migrating to local draining lymph nodes (Bellingan et al., 1996; Gautier et al., 2013). However, under persistent inflammation, some of the recruited SPMs eventually differentiate into F4/80^high^ MHC II^low^ cells, which corresponds to the intermediate subtypes identified in this study (Bain et al., 2016; Yona et al., 2013). Consistently, by constructing a continuous developmental trajectory using cells of DC1, SPMs, and IMS, we reported that ELL-induced inflammation promoted the differentiation of SPMs from DC1 by increasing the expression of genes necessary for this process. Moreover, our analysis strongly suggests that *Retnla* is a potential key driver for this process. In support of this, reduced expression of *Retnla* by niclosamide suppressed this early differentiation process and led to reduced populations of both recruited Ly6C+ monocytes and more differentiated FRβ+ IMS, indicating that niclosamide is able to improve persistent or chronic information that is developed by lesion establishment.

Severe inflammation results in the ablation of embryo-derived RLMs and increases the replenishment of RLMs by monocyte-derived macrophages (Louwe et al., 2021). The two populations of IML1 and IML2 are in the transition to acquiring their long-term residence as embryo-derived RLMs. By constructing a maturation process of IML1 and IML2, we found that ELL decreased the expression of necessary genes such as *Cfb* to promote, while niclosamide enhanced most of these gene expressions to reverse this replenishment process. Moreover, both the expression of *Timd4* and the cell number of TIM4+ RLMs were decreased by ELL and increased by niclosamide. Therefore, niclosamide prohibited the replenishment of resident macrophages by both suppressing the maturation of monocyte-derived macrophages and preserving the population of embryo-derived RLMs. As the monocyte-derived RLMs were reported to have different functions from original RLMs, enhanced replenishment of resident macrophages may have long-term disruptions to the peritoneal niche of endometriosis (Louwe et al., 2021).

CXCL13 expression in TIM4+ RLMs plays an essential role in the maintenance and recruitment of B1 cells from circulation (Beattie et al., 2016; Zeng et al., 2018). However, the expression of CXCL13 is deficient in monocyte-derived RLMs, which leads to disruption of B1 cell homeostasis under inflammation (Bain et al., 2020; Louwe et al., 2021). We found that CXCL13 was actually expressed in all LPMs (IML1, IML2, and RLMs). But, consistently, we found that ELL decreased the cell number of TIM4+ RLMs, and the expression of CXCL13 in macrophages. Niclosamide promoted the expression of CXCL13 and rescued the communications of macrophages to B1 cells. In addition, intercellular interaction analysis also suggested that the three “large” types of macrophage populations might also regulate the population of B2 cells by the *C3* expression. Treatment of niclosamide enhanced the expression of *C3*. Therefore, ELL might disrupt the communications between LPMs and B1 and B2 cells, and niclosamide could rescue this communication through its regulations on macrophages.

In summary, the heterogeneity and developmental characteristics of peritoneal macrophages were extensively explored this study using a mouse model of endometriosis. ELL enhanced the process of early differentiation and maturation of monocyte-derived macrophages while reducing the maintenance of embryo-derived RLMs. The increased replenishment of resident macrophages by monocyte-derived ones further disrupts the homeostasis in the peritoneal cavity and affects the recruitment and functional activities of B cells. Niclosamide stepwise reverses the dynamic progression of recruited macrophages and preserves the population of embryo-derived RLMs, hence tuning the perturbed peritoneal microenvironment in endometriosis back to normal. Therefore, macrophages could be a direct target of niclosamide, and niclosamide could be a new therapy to recuse perturbed peritoneal microenvironment that contributes to chronic inflammation, lesion growth, and progression, neuroangiogenesis, and endometriosis-associate pain.

## Materials and methods

### Animals and a mouse model of endometriosis

All procedures were performed in accordance with the guidelines approved by the Institutional Animal Care and Use Committee of the Washington State University (Protocol # 6751). C57BL/6J mice were purchased from the Jackson Laboratory. Endometriosis-like lesions (ELL) were induced by inoculating syngeneic menstrual-like endometrial fragments from donor mice into the peritoneal cavity of recipient mice, as described previously (Greaves et al., 2014; Shi et al., 2021). Briefly, ovariectomized donor mice were primed with estradiol-17β (E2) and progesterone and induced decidualization by injecting sesame oil into uterine horns to produce a “menses-like” event. Then, decidualized endometrial tissues were scarped from myometrium, minced, and injected i.p. into ovariectomized and E2-primed recipient mice under anesthesia (50 mg tissue in 0.2 mL PBS per recipient). Sham mice were ovariectomized and E2-primed and injected with 0.2 mL PBS into the peritoneal cavity. Three weeks after the induction of ELL or sham, mice in the groups of sham, ELL, and ELL_N were orally administrated with vehicle or niclosamide (200 mg/kg/day) for a total of 3 weeks (Fig. 1A), as described previously (Shi et al., 2021). At 6 weeks following ELL induction, mice were euthanized, and peritoneal cells were collected for further analysis.

### Preparation of peritoneal immune cells for single-cell RNA sequencing

Peritoneal immune cells were isolated and collected from the peritoneal fluid following our established method (Shi et al., 2021). After removing red blood cells by lysis, remaining cell mixtures were used for cDNA library constructions following the manufacturer’s protocol (10X Genomics, Inc.) of the Chromium Single Cell 3’ Library & Gel Bead Kit V3 (Zhao et al., 2021). All samples were multiplexed together and sequenced across one single lane of an Illumina NovaSeq 6000 S4.

### Single-cell data processing and analysis

Raw data in FASTQ format were pre-processed with Cell Ranger V3.1.0 (10x Genomics) mapping to the mouse GRCm38/mm10 transcriptome to generate gene-cell matrices. A total of 13,859 cells from all three libraries were integrated into R using the Seurat package (V4.0.4) (Stuart et al., 2019). To filter out doublets and low-quality cells, criteria of 750,000 unique molecular identifiers (UMIs) and 500 genes per cell were set. In addition, cells with over 20% expression of mitochondrial genes were excluded for downstream analysis. Modified multivariate Pearson’s RV correlations for each set of treatment replicates were calculated using the package of MatrixCorrelation (v0.9.2) (Smilde et al., 2009). The following correlations showed consistent sampling between libraries of treatments: ELL and sham = 0.962, ELL_N and sham = 0.979, ELL and ELL_N = 0.985. The “sctransform” function was then applied to normalize the remaining dataset with regression of mitochondria mapping percentage (Hafemeister and Satija, 2019). Dimensionality reduction was performed on identified variable genes by principal component analyses (PCA). The top 66 dimensions were selected for clustering (“resolution” set to 0.5) and uniform manifold approximation and projection (UMAP) visualization. Specific gene markers for each cluster were identified using the “FindAllMarkers” function. Differential gene expression between treatments was analyzed using the “wilcox” test in the “FindMarkers” function with Bonferroni adjusted *p* value < 0.05 showing significant differences. Gene Ontology (GO) and Gene Set Enrichment Analysis (GSEA) was performed with the R package, clusterProfiler V3.18.0, using all detected genes from the entire scRNA-seq library as background (Yu et al., 2012). Terms were enriched with the nominal *p* value < 0.05 and false discovery rate (FDR) (*q* value) < 0.05.

### An interactive web tool to share scRNA-seq data of peritoneal immune cells

Single-cell transcriptomic analysis has provided an unprecedented high resolution of peritoneal immune cells including different subtypes of B cells, macrophages, and T cells. To share our data with other researchers, we have created a cloud-based web tool for easy gene searches, which does not require complicated computer programming skills (Thompson et al., 2021). The webpage for this tool is: https://kanakohayashilab.org/hayashi/en/mouse/peritoneal.immune.cells/

### Single-cell trajectory analysis

The biological processes of “small” macrophage recruitment and “large” macrophage maturation were revealed by the trajectory analysis. Cells from clusters of DC1, SPMs, and IMS or IML1, IML2, and RLMs were computationally selected in Seurat, and the two data matrices were imported, processed, and pseudo-ordered using the package of Monocle 3 in R following the standard pipeline (Qiu et al., 2017).

### *In Silico* Knockout Analysis

Functional analysis of *Retnla*, *Cfb,* and *Timd4* was conducted using the R package of scTenifoldKnk V1.0.1 (Osorio et al., 2022). A single-cell gene regulatory network (scGRN) was conducted using our scRNA-seq data from the sham group. Then the expression of *Retnla*, *Cfb,* and *Timd4* was set to zero from the constructed scGRN to build their own corresponding “pseudo-knockout” scGRN. Perturbed genes by this virtual knockout were quantified by comparison of the “pseudo-knockout” scGRN to the original scGRN. Those significantly affected genes were used for GO analysis to show changes in biological processes caused by *in silico* knockout.

### Intercellular Communication Analysis

Gene expression data of Seurat objects were used as input to model the probability of intercellular interactions between B cells and macrophages using the R package of CellChat V1.0.0 (Jin et al., 2021). The known database of interactions (CellChat.DB.mouse) between ligands, receptors, and cofactors was used as the reference.

### Re-analysis of one public-available single-cell dataset of peritoneal macrophages

A single-cell transcriptomic dataset of peritoneal macrophages from 19-weeks old female mice is pubic available (Bain et al., 2020). These cells were selected based on the expression of CD11b with the removal of granulocytes and B1 B cells using flow cytometry. The raw data were downloaded from NCBI GEO (GSM4151331), pre­processed with Cell Ranger V3.1.0, and re-analyzed in the R package, Seurat V4.0.4. For quality control, cells with the expression of fewer than 300 genes or over 5000 genes, and over 5% mitochondrial genes were excluded, resulting in a total of 4287 out of 4702 cells for downstream analysis. Following the standard pipeline of Seurat, the raw counts were normalized using a global-scaling normalization method (“LogNormalize”). Using the “ScaleData” function, the normalized dataset was further scaled for dimensional reduction. Based on the principal component analyses (PCA), the top 34 dimensions were selected for clustering and UMAP graphing. Trajectory analysis was performed on the cells computationally selected from the “small” or “large” macrophage lineages as described above using the R package of Monocle 3.

### Flow Cytometry

Peritoneal cells were harvested and used for analyzing immune cell profiles by flow cytometry. Briefly, the peritoneal lavages were centrifuged to collect peritoneal exudate cells. After lysing red blood cells by 1x RBC Lysis Buffer (BioLegend), an equal number of cells from each group were incubated at room temperature for 20 minutes with Zombie Aqua™ Fixable Viability dye (BioLegend) and blocked on ice for 20 minutes with FcBlock anti-CD16/CD32 (Thermo Fisher). Then cells were stained with fluorochrome-conjugated monoclonal antibodies (Table S1) for 1 hour. Samples were acquired with the Attune NxT Acoustic Focusing Cytometer using Attune NxT software (Invitrogen), and data were analyzed with FlowJo v10.4. For analysis, only singlets (determined by forward scatter height vs. area) and live cells (Zombie Aqua negative) were used.

### Quantitative Real-time PCR Analyses (qPCR)

Total RNA was isolated using TRIzol reagent (Sigma #T9424), and cDNA templates were synthesized from 1µg of purified RNA using the High-Capacity cDNA Reverse Transcription Kit (Thermo Fisher) (Shi et al., 2021; Zhao et al., 2020). qPCR was performed using a CFX RT-PCR detection system (Bio-Rad), and relative gene expression was evaluated by SYBR Green (Bio-Rad #1725274) incorporation. *Rpl19* was used as the reference gene to normalize mRNA expression levels. Data were analyzed using the 2^-ΔΔCt^ method. Primer sequences were provided in Table S2.

### Statistical Analysis

For single-cell transcriptomic sequencing, in each treatment, a total of 3 mice were used for sample preparation. Pre-processing of raw sequencing data including transformation, normalization, and quality control were described above. For the analysis of differential gene expression, the default “wilcox” test was performed using the R package Seurat (v4.0.4). For RT-qPCR of peritoneal immune cells, six mice from each treatment were used for RNA extraction (n=6). For flow cytometry, cells from three mice were pooled as one sample and a total of 15 mice were used for each group of treatments (n=5). For comparisons between three groups of treatments, one-way ANOVA followed by Tukey’s multiple comparisons was used. Data were analyzed with GraphPad Prism (version 9) and presented as means ± SEM. Statistical differences were indicated as \**p* < 0.05, \*\**p* < 0.01, \*\**p* < 0.001.

### Data availability

The data that support the findings of this study are openly available in the GEO database at NCBI, reference number GSE147024.

## Acknowledgments

The study was supported by the National Institutes of Health (NIH), Eunice Kennedy Shriver National Institute of Child Health & Human Development (NICHD) R01HD104619 (to K. Hayashi). The graphical abstract was created using BioRender.com.

## Author contributions

M. Shi and K. Hayashi designed experiments. L. Zhao and M. Shi performed experiments and analyzed data. S. Winuthayanon assisted with the webtool constructions and technical support with data analysis. J. A. MacLean and K. Hayashi assisted in experiments and provided critical feedback on the manuscript. L. Zhao wrote the paper. All authors read, edited, and approved the manuscript.

## Online supplementary material

**Figure S1** shows the characteristic gene expression of immune cells at the single-cell level. **Figure S2** shows characteristic profiles of each macrophage subpopulation. **Figure S3** shows the re-analysis of a single-cell transcriptomic dataset of peritoneal macrophages in female mice in a normal physiological state from a public resource. **Figure S4** shows the construction of a trajectory path for the differentiation of recruited macrophages (data from a public resource). **Figure S5** shows the construction of a trajectory path for “large” macrophages. **Table S1** shows enriched GO terms of biological processes in each macrophage subpopulation. **Table S2** shows enriched GSEA terms of biological processes in macrophages between treatments. **Table S3** shows genes and GO biological processes affected by *in silico* knockout of *Retnla*, *Cfb,* and *Timd4*. **Table S4** shows enriched GSEA terms of biological processes in B cells between treatments. **Table S5** shows antibodies and reagents for Flow Cytometry. **Table S6** shows primer information used for RT-qPCR.

## Supplemental Figure Legends

**Figure S1 Characteristics of immune cells at the single-cell level.** (A) Dot plot showing one selected lineage-specific marker gene expression for each cluster. (B) UMAP distribution of cell populations within three groups. (C) Characteristic expression of marker genes for macrophage subpopulations. Sham, sham control; ELL, endometriosis-like lesions; ELL_N, niclosamide administration to ELL-induced mouse; DC1, dendritic cells 1; SPMs, “small” peritoneal macrophages; IMS, intermediate “small” macrophages; IML1, intermediate “large” macrophages subtype 1; IML2, intermediate “large” macrophages subtype 2; RLMs, resident “large” macrophages; PMs, proliferating macrophages; B1a, B1a cells; B1b, B1b cells; B2, B2 cells; GC-Bs, germinal-center B cells; PB-Bs, plasma blast B cells; Cd4+ Ts, Cd4+ T cells; Cd8+ Ts, Cd8+ T cells; Cd4-/Cd8-Ts, Cd4­/Cd8-T cells; Blast Ts, blast T cells; DC2, dendritic cells 2; MCs, Mast cells; NTPs, neutrophils.

**Figure S2 Enriched GO terms of biological processes by the top 100 expressed genes within each macrophage subpopulation to show their characteristic profiles.**

DC1, dendritic cells 1; SPMs, “small” peritoneal macrophages; IMS, intermediate “small” macrophages; IML1, intermediate “large” macrophages sybtype 1; IML2, intermediate “large” macrophages subtype 2; RLMs, resident “large” macrophages.

**Figure S3 Re-analysis of a single-cell transcriptomic dataset of peritoneal macrophages in female mice in a normal physiological state from a public resource.**

(A) UMAP visualization of macrophage populations. B) Gene markers used for macrophage subpopulation clustering. (C) Feature plot showing the distribution of genes. DC, dendritic cells; SPM, “small” peritoneal macrophages; IMS, intermediate “small” macrophages; LPM1-4, intermediate “large” macrophages 1-4; PMs, proliferating macrophages.

**Figure S4 Reconstruction of a trajectory path for the differentiation of recruited macrophages (data from a public resource)**

(A) UMAP showing the selected cells (DC, SPM, and IMS) and the trajectory path built within them. (B) Dynamic expression of genes along this trajectory path. DC, dendritic cells; SPM, “small” peritoneal macrophages; IMS, intermediate “small” macrophages.

**Figure S5 Reconstruction of a trajectory path for “large” macrophages**

(A) UMAP showing the selected cells (LPM1-4) and the trajectory path built within them. (B) Dynamic expression of genes along this trajectory path. LPM1-4, intermediate “large” macrophages 1-4.

