## Supplementary figures and images for "Niclosamide targets macrophages to rescue the disrupted peritoneal homeostasis in endometriosis"

### Fig S1

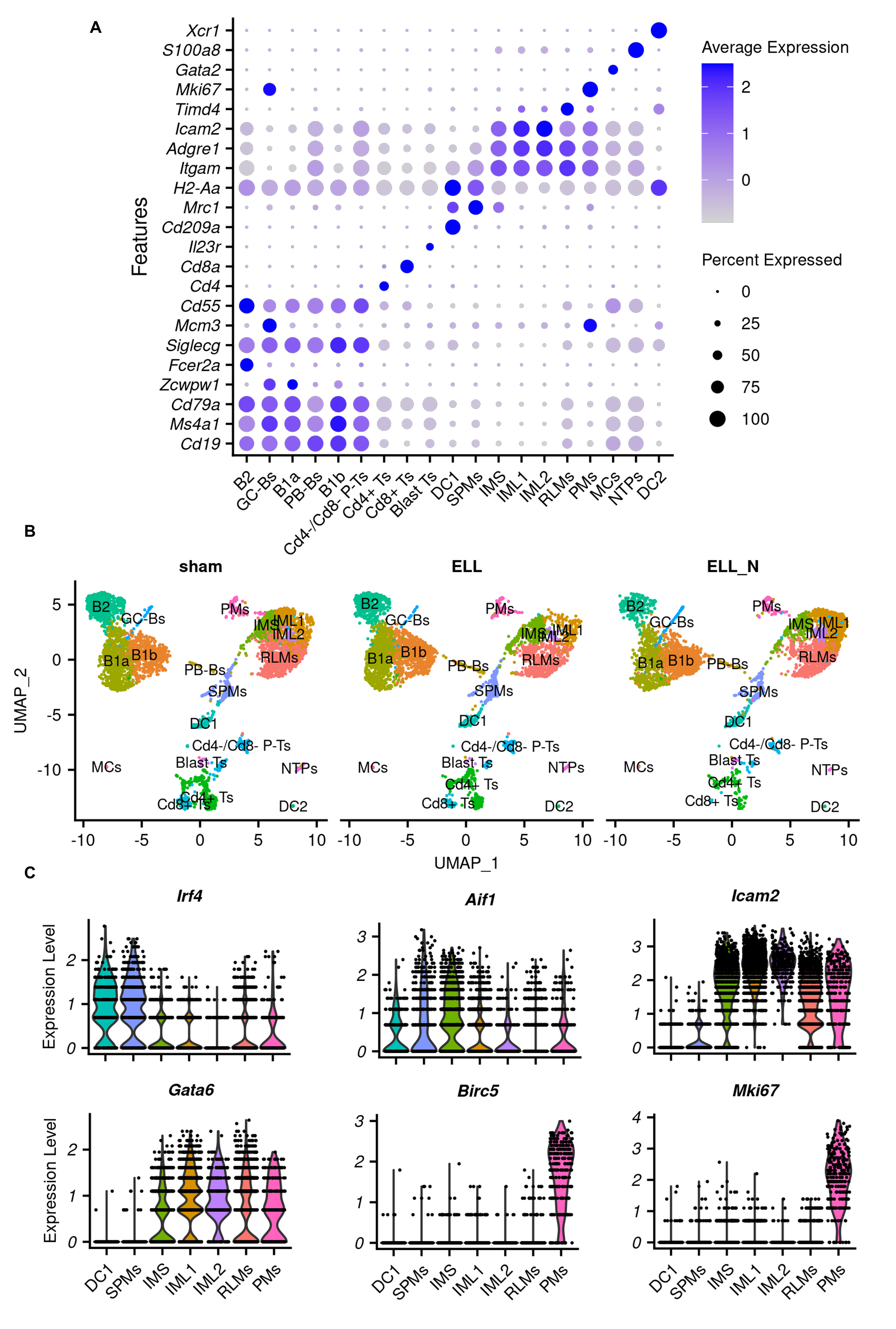

### Fig S2

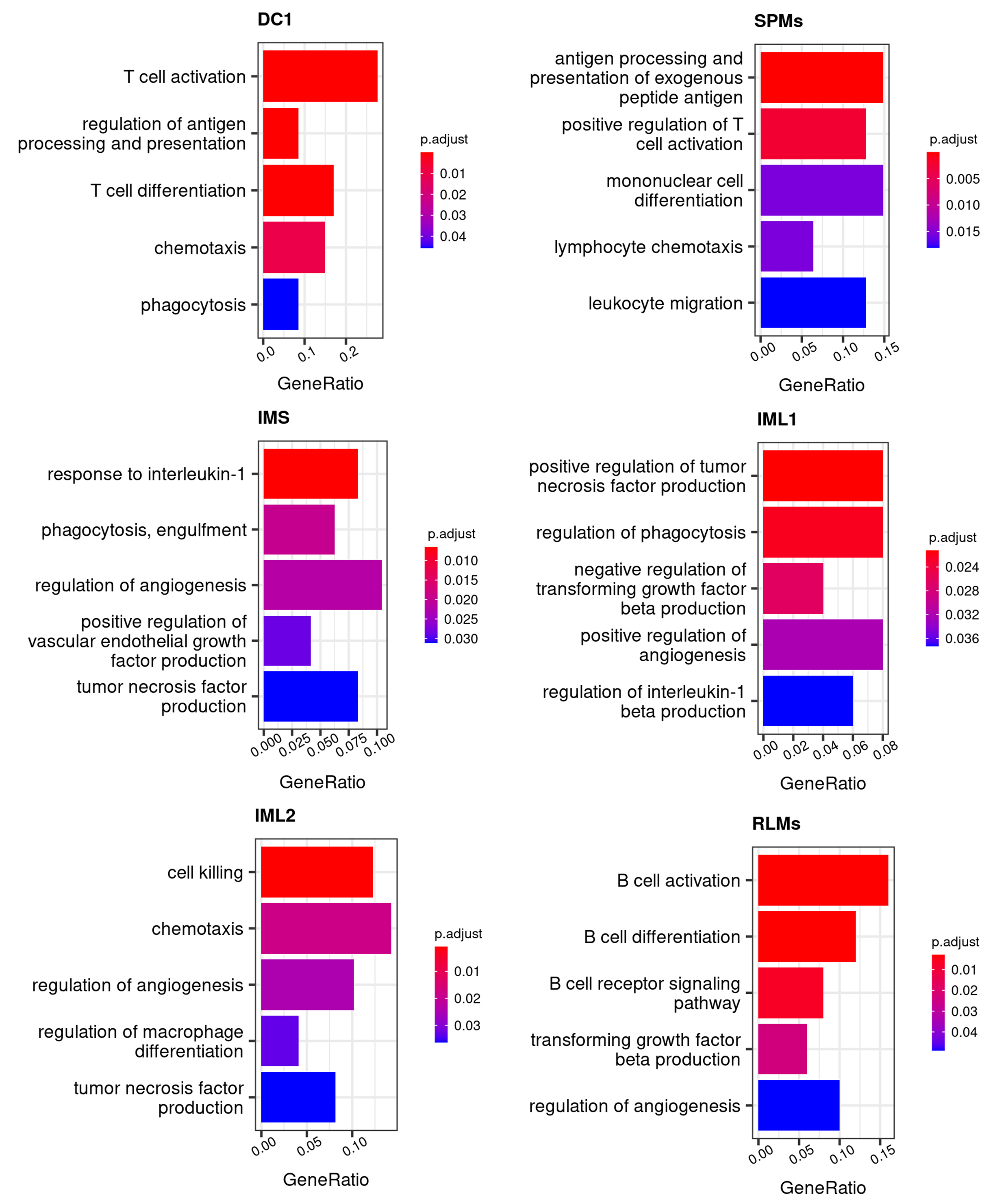

### Fig S3

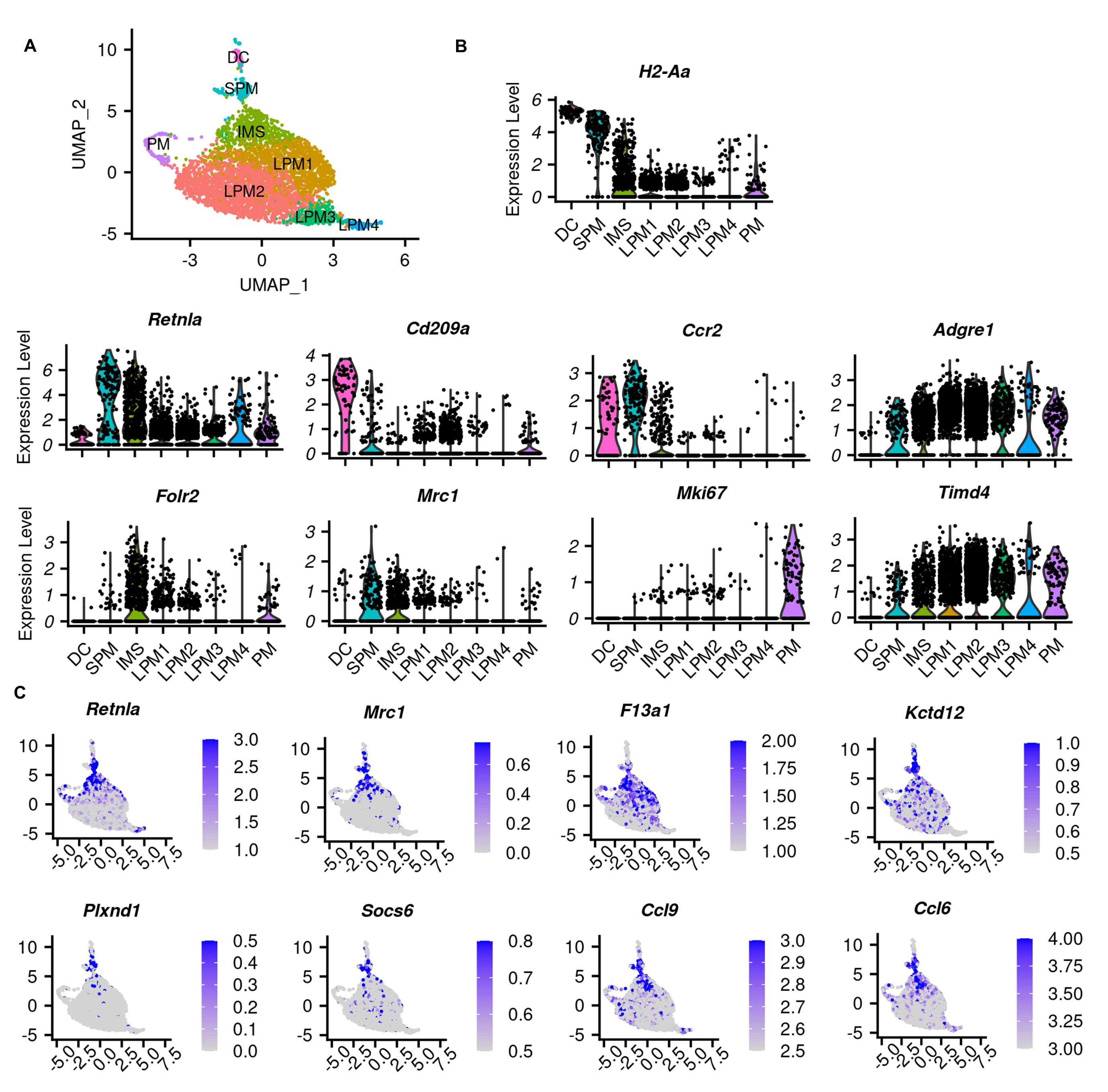

### Fig S4

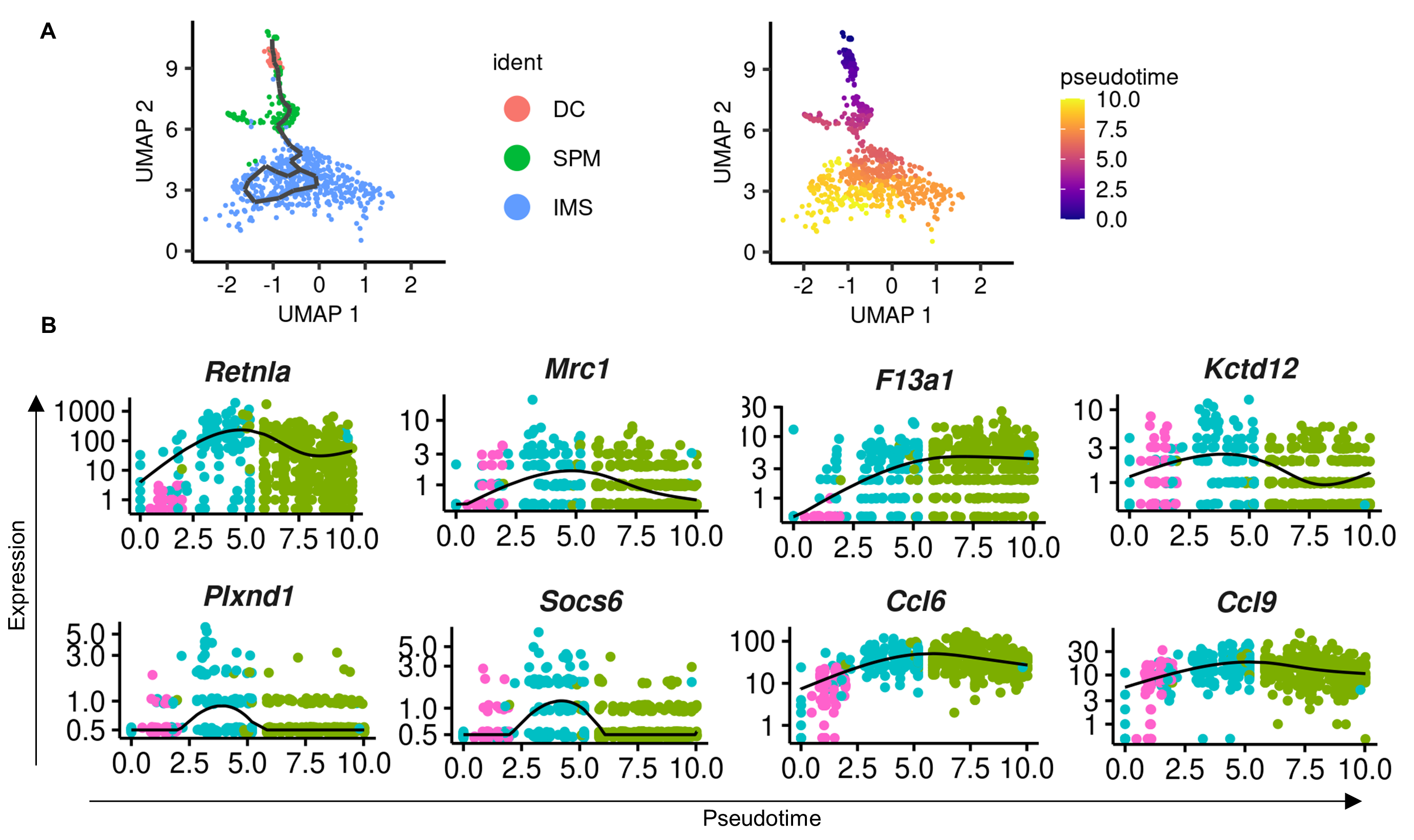

### Fig S5

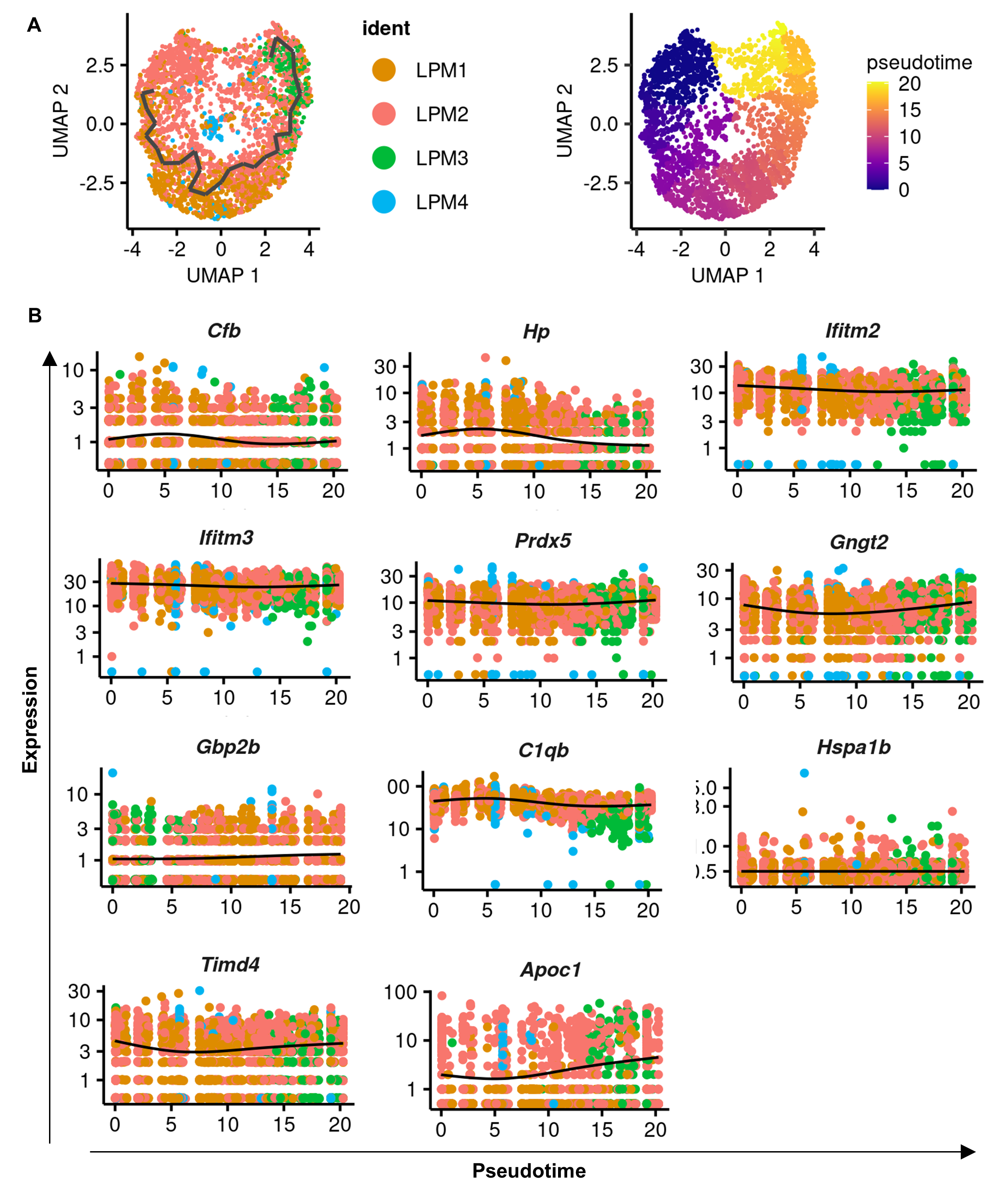
